# An Automated Model Annotation System (AMAS) for SBML Models

**DOI:** 10.1101/2023.07.19.549722

**Authors:** Woosub Shin, John H. Gennari, Joseph L. Hellerstein, Herbert M. Sauro

## Abstract

**Motivation:** Annotations of biochemical models provide details of chemical species, documentation of chemical reactions, and other essential information. Unfortunately, the vast majority of biochemical models have few, if any, annotations, or the annotations provide insufficient detail to understand the limitations of the model. The quality and quantity of annotations can be improved by developing tools that recommend annotations. For example, recommender tools have been developed for annotations of genes. Although annotating genes is conceptually similar to annotating biochemical models, there are important technical differences that make it difficult to directly apply this prior work.

**Results:** We present AMAS, a system that predicts annotations for elements of models represented in the Systems Biology Markup Language (SBML) community standard. We provide a general framework for predicting model annotations for a query element based on a database of annotated reference elements and a match score function that calculates the similarity between the query element and reference elements. The framework is instantiated to specific element types (e.g., species, reactions) by specifying the reference database (e.g., ChEBI for species) and the match score function (e.g., string similarity). We analyze the computational efficiency and prediction quality of AMAS for species and reactions in BiGG and BioModels and find that it has sub-second response times and accuracy between 80% and 95% depending on specifics of what is predicted. We have incorporated AMAS into an open-source, pip-installable Python package that can run as a command-line tool that predicts and adds annotations to species and reactions to an SBML model.

**Availability:** Our project is hosted at https://github.com/sys-bio/AMAS, where we provide examples, documentation, and source code files. Our source code is licensed under the MIT open-source license.

**Supplementary information:** Supplementary data are available online.

## 1. Introduction

In systems biology, annotations are metadata that can describe models and model elements. For example, annotations can provide detailed information about the chemical species that may participate in reactions in a model. These annotations leverage canonical, readily available knowledge resources such as the Gene Ontology and ChEBI (Degtyarenko et al. 2008; Ashburner et al. 2000). Annotations on a biosimulation model provide contextual information that can be used by researchers and systems for advanced search capabilities and to determine suitability for reuse (Cowan, Mendes, and Blinov 2019; Neal et al. 2019; Schulz et al. 2011). For example, researchers have demonstrated that complex biological models can be constructed by combining simpler models if appropriate annotations are present (Krause et al. 2010; Snoep et al. 2006).

A number of tools are based on the assumption of high-quality and pervasive model annotations. semanticSBML by Krause *et al*. merges models that have annotations in common for model elements (Krause et al. 2010). ModelBricks builds on the idea of high-quality annotations of model elements to construct reusable model modules that can be easily incorporated into new models (Cowan, Mendes, and Blinov 2019). Finally, Sarwar *et al*. demonstrate the ability of annotations to support specialized and detailed search capabilities (Sarwar et al. 2019). These examples illustrate the benefits that can be derived by having well-annotated models.

Annotating model elements is a time-consuming and knowledge-intensive task. For chemical species, choosing an appropriate annotation from the ChEBI knowledge resource often requires a detailed examination of a large number of similar molecules. For example, the name “glucose” is associated with approximately 1,000 ChEBI entries. The effort to annotate an entire model is the effort to annotate a single model element multiplied by the number of species, reactions, and other elements in the model. Because this is a substantial effort, models are typically not well annotated. As we described in our earlier analysis (Shin et al. 2021), about half of the models in the BioModels collection have fewer than half of their species and reactions annotated. Even for models that have some annotations, there are significant gaps that can impair the understanding of models and their results. A common reason for *not* annotating model elements is that doing so is a painstakingly difficult manual effort.

The need for high-quality annotations is well recognized in other areas of biological modeling. For example, extensive tooling (e.g., Manning, Raghavan, and Schütze 2008) has been developed for gene annotations, and more broadly, for annotation of common biological polymers (e.g., DNA, RNA, proteins). This has been successful because functionally equivalent biological polymers are readily identified by the similarities in their atomic (or residue) sequences. Tools such as BLAST and HMMER provide high-quality and efficient algorithms for calculating match scores (Finn, Clements, and Eddy 2011; McGinnis and Madden 2004). An unannotated molecule is annotated by searching a database of annotated molecules to find molecules with a high match score with the unannotated molecule. There are several databases available for analyzing biological polymers, including RefSeq and GeneBank for genes and pFam and uniProt for proteins (Leray et al. 2019; Mistry et al. 2021; Pruitt, Tatusova, and Maglott 2007; Consortium 2014).

The foregoing capabilities have been impactful. For example, ModelSEED, a web resource for genome-scale reconstruction and analysis, obtains the annotation of assembled genome sequence using RAST, an automated annotation service for archaeal and bacterial genomes (Aziz et al. 2008; Henry et al. 2010; Mendoza et al. 2019). Other examples include merlin (Dias et al. 2015; Mendoza et al. 2019) and Architect (Nursimulu, Moses, and Parkinson 2022).

However, can these techniques be applied to annotating elements of SBML models? From the foregoing, there are two requirements: (1) there must be a reference database of annotated model elements; and (2) there must be a way to find elements in the reference database that are similar to an unannotated element in a model. The first condition is satisfied for chemical species (e.g., ChEBI) and reactions (e.g., Rhea). However, the second condition is more difficult.

The challenge is that there is no simple way to apply sequence similarity to many model elements, especially species and reactions. Genes can be described by a long sequence of a small alphabet – {G, A, T, C}. This tends to make sequence similarity effective for identifying similar genes. In contrast, it is non-trivial to apply sequence similarity to chemical species. We cannot use the similarity of chemical structures because the chemical structure of an unannotated species is not known. Further, sequence similarity is problematic for reactions since the alphabet consists of thousands of different molecules that could be reaction participants.

We introduce AMAS, an Automated Model Annotation System that predicts annotations for chemical species and reactions in SBML models. The core of our approach is the development of appropriate similarity measures, which we refer to as **match scores**. We do this by leveraging two properties of SBML models. First, elements are typed so that we know if a symbol is a chemical species, a reaction, or some other component of the model. Second, all SBML elements have an identifier, and some elements have user-provided display names as well. Finally, SBML reactions indicate which chemical species participate as reactants or products. As we describe below, our methods also leverage ChEBI for reference information about chemical species (Degtyarenko et al. 2008), and Rhea for information about reactions (Alcántara et al. 2012). Our match scores for chemical species are based on element identifiers and/or display names. Our match scores for reactions are quantified in terms of the similarity of the chemical species that participate in that reaction.

Others have aimed to automate annotations. For example, SBMLSqueezer 2 predicts reaction annotations based on species participants (Dräger et al. 2015). However, this approach requires that the chemical species have annotations, whereas we first predict un-annotated species, and then use this information to optimize our predicated annotations for the reactions. Similarly, Leonidou *e*t al. describe an approach (Leonidou et al. 2023) that relies on having correctly annotated species to permit the analysis of reaction types (e.g., transport reactions). Although the paper demonstrates effectiveness in terms of providing more precise annotations for BiGG models, its scope is limited to Systems Biology Ontology (Courtot et al. 2011) annotations of reactions. In particular, it does not address the absence of annotations of species, a major consideration in the BioModels repository.

The contributions of this work are: (a) construction of an accurate and computationally efficient way to predict annotations for species and reactions in SBML models, (b) the implementation of a tool (AMAS) that recommends annotations and can add those annotations into an SBML model, and (c) an evaluation of these methods and this tool with models from the BiGG and BioModels collections. AMAS is an open-source, pip-installable Python package.

## 2. Methods

Fig. 1 displays an overview of our method. Beginning with the left side of Fig. 1, a **query element** is an element whose annotation is to be predicted. The **Element Filter** determines if AMAS can predict any annotations for a query element. For example, some models name species in a way that provides no useful information for predicting annotations (e.g., S1, S2, …). The Element Filter rejects these query elements. We assume there are **reference elements** whose annotations are known, and these elements reside in a **reference database**. A **Match Score Function** takes as arguments a query element and a reference element; it returns a **match score**, a number in [0, 1]. A match score of 0 indicates that the two elements are maximally *dissimilar*; a score of 1 indicates maximal *similarity*. The **Match Score Selector** chooses a subset of the annotations as the **prediction set** of annotations based on the match score of the reference element associated with the annotation. Query elements rejected by the element filter have an empty prediction set.

**Fig. 1.**
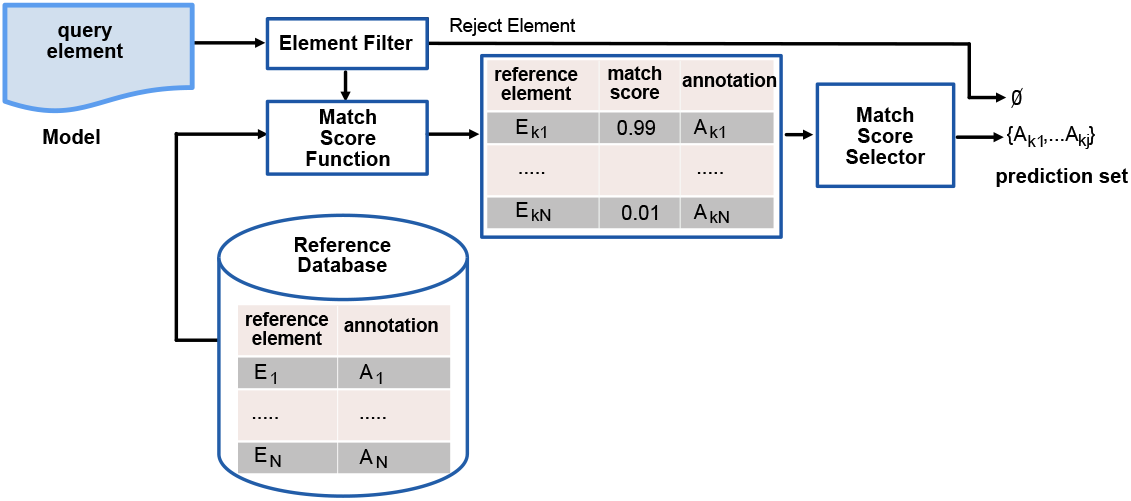
Summary of AMAS annotation prediction. Annotation prediction is done for a query element by choosing the annotations of reference elements (with known annotations) that have the highest match scores as calculated by a Match Score Function. The Element Filter determines if there is sufficient information about a query element to make any prediction.

The Match Score Selector can use two different **match score selection criteria (MSSC)**:

- **MSSC Top:** Predicted annotations are those associated with the reference element with the largest match score above a user-specified minimum value called the **match score cutoff**. If multiple reference elements have the same score, then their annotations are predicted as well.
- **MSSC Above:** Predicted annotations are all reference elements whose match score exceeds the match score cutoff.

Both MSSCs are useful, as we detail below.

AMAS may produce an empty prediction set. This occurs if the query element is rejected by the Element Filter. It also occurs if MSSC Cutoff is used and the largest match score for the query element is smaller than the match score cutoff.

We use three metrics to assess the quality of AMAS predictions. **Nonempty** is the percentage of annotation predictions where there is at least one annotation in the prediction set. **Accuracy** is the probability that a correct annotation of a query element is in the prediction set if the prediction set is not empty. If the query element has multiple correct annotations, then accuracy is the probability of at least one correct annotation being in the prediction set. Last, we consider the size of the prediction set: **Exactness** quantifies how specific the prediction is: the inverse of the size of the prediction set. Nonempty, accuracy, and exactness are in [0, 1] with 1 being the ideal value.

We have developed an application that uses AMAS predictions to recommend annotations for species and reactions for SBML models. The approach is: (a) predict annotations for the query element; (b) if MSSC Top is used, display the annotations for the reference element(s) with the largest match score above a user-specified threshold; (c) if MSSC Above is used, display the annotations for the reference elements whose match score exceeds the threshold. In both cases, it can be valuable to the user to see the match scores associated with the annotations (via the reference element for that annotation).

### 2.1 Predicting Species Annotations

AMAS calculates the similarity between two species based on the similarity of strings associated with the two species. For the query species, the preferred strings is the SBML display name if it exists (since this tends to better reflect the nature of the species). If the display name is absent, we use the SBML element identifier. The Element Filter for predicting species annotations is the length of the string for the element; our experience is that a length of at least 3 or 4 characters provides good results. In our current implementation, the reference species is a term in the ChEBI database. Each ChEBI term has a list of frequently used synonyms, and these synonyms are the strings associated with the reference species. For example, the synonyms for CHEBI:17634 include D(+)-glucose, dextrose, and grape sugar.

String similarity is conceptually simple, but the details can be a bit more complex. Strings may have different lengths. One string may be a sub-string of the other. Letters may be inserted, deleted, and/or transposed. Given these considerations, what is a meaningful score that quantifies the similarity of two strings? Approaches such as the Needleman–Wunsch algorithm (Needleman and Wunsch 1970) assume much longer strings, and work best with a small alphabet and non-binary scoring for character mismatches.

These computational concerns led us to calculate string similarities using cosine similarity, a technique used in Natural Language Processing (Han, Kamber, Pei, et al. 2012). A string is represented as a vector in a 36-dimensional space: one dimension for each letter of the alphabet and for the numerals 0 through 9. The vector representation of a string has a “1” as the coordinate of a dimension in which that character is present in the string; otherwise, the coordinate is zero. For example, the vector for “ATP” has a “1” as the coordinate for the “a”, “t”, and “p” dimensions of the vector space, and a “0” for all other coordinates. We quantify the similarity of two strings as the cosine of the angle between their vector representations. Because of the structure of the space, the cosine is in [0, 1] (since the dot product can never be negative because all coordinates are non-negative). A value of 1 is obtained when the angle is 0. This means that the vectors coincide, and so the query string and the synonym are ideally similar. The match score between a query species *S*^*q*^ and a reference species *S*^*r*^ is the maximum cosine distance between the (string) name *N* ^*q*^ for *S*^*q*^ and the names of the synonyms 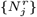 for *S*^*r*^:

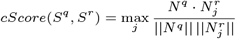

where 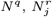 are vector representations of species names and ||*N* || is the length of the string *N* in the vector space.

One concern with *cScore* is that the encoding is lossy in that we cannot recover the original string from its vector representation. More elaborate encodings can reduce this problem. However, we found that the simple binary encoding produced good results for predicting annotations. Another concern is that several ChEBI terms can share a synonym, and so the accuracy of predictions could be impaired if the two or more reference species share a synonym that has the largest cosine distance with a query species. This said, our evaluations in Section 3.1 suggest that AMAS produces reasonably accurate predictions.

### 2.2. Predicting Reaction Annotations

This section describes how we predict reaction annotations using the framework presented in Fig. 1. As with predicting species annotations, we start by constructing a match score that quantifies how close a query reaction is to a reference reaction. Analogous to predicting species annotations, reference reactions are in a database and have associated annotations. We use Rhea (Alcántara et al. 2012) for the database of reference reactions.

Let *rScore*(*R*^*q*^, *R*^*r*^) be the match score between the query reaction *R*^*q*^ and the reference reaction *R*^*r*^. We calculate *rScore* based on the similarity between the participants of the query reaction and the participants of reference reactions.

The first step in calculating *rScore* is to simplify the chemical formulas of reaction participants. We do this by omitting all hydrogen atoms. There is one exception to this rule. *H*_*n*_ has the simplified formula of *H*. Using simplified chemical formulas helps with handling chemical variants created by protonation and deprotonation, and it reduces computational complexity (by having fewer chemical formulas).

As with species annotations, we represent reactions as binary valued vectors. The dimension of the vector space are the simplified chemical formulas of species in the ChEBI database. The *i*-th coordinate in a reaction vector is 1 if there is at least one participant with that simplified chemical formula; otherwise, the coordinate is 0. If a species has more than one chemical formula, then there may be more than one non-zero coordinate associated with that species.

Let *R*^*q*^ be the vector representation of the query reaction and *R*^*r*^ be the representation of a reference reaction. *R*^*q*^ · *R*^*r*^ is the dot product of these vectors, the sum of the product of the coordinates of the two vectors. Note that this is the number of simplified chemical species that are common to the two reactions. The reaction match score is proportional to *R*^*q*^ · *R*^*r*^.

We have found that the AMAS match score works best if an appropriate normalization constant is applied to *R*^*q*^ · *R*^*r*^. The normalization constant *D* is a function that considers *all* reference reactions that have the largest dot product with *R*^*q*^. We denote this largest dot product by *M* ^*q*^, and indicate distinct reference reactions by 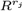. The resulting set is 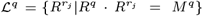. The normalization constant is the smallest number of participants for reactions in ℒ^*q*^. That is, 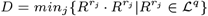. The AMAS reaction match score is:

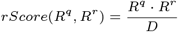

Note that *D* does not influence the ranking of match scores between the query reaction and the reference reactions. Rather, *D* affects the *value* of the match score, and this has implications for the prediction set: annotations are included in this set only if their scores exceed the match score cutoff value.

As an example, consider the query reaction *ATP* → *ADP*, a simplified description of ATP hydrolysis. *R*^*q*^ for this reaction has a 1 in the coordinate for *ATP* and a 1 in the coordinate for *ADP*. We calculate *rScore*(*R*^*q*^, *R*^*r*^) for the reference reaction RHEA:13065, which has the participants *ATP, H*_2_*O, ADP, H*^+^, and 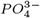 (phosphate). Note that *R*^*q*^ ·*R*^*r*^ = 2 = *M* ^*q*^ (since the query reaction only has two participants). For this example, we assume that 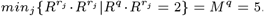. Hence, *D* = 5. So, 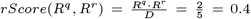. Section S5 contains more details on the calculation of *rScore*.

### 2.3. Optimizing Annotation Predictions

Once reaction annotations have been predicted, there is an opportunity to improve the prediction of species annotations. In effect, we can use information about participation in a known reaction to make better, more complete guesses about species annotation.

Fig. 2 illustrates the foregoing. On the left is a query reaction *R*_*ACKr*, and on the right is the reference reaction with the largest match score, RHEA:11352. The figure displays how participants of the query reaction are paired with participants of the reference reaction. We see that in three of the four pairings, the annotations are same or are “matched”. However, the species *M atp c* is paired with a species whose annotation differs from that assigned to *M*_*atp*_*c*. So, if RHEA:11352 is the correct annotation for the query reaction, then we should change the annotation of *M*_*atp*_*c* to CHEBI:20616 since this is the annotation its paired species in the reference reaction.

**Fig. 2.**
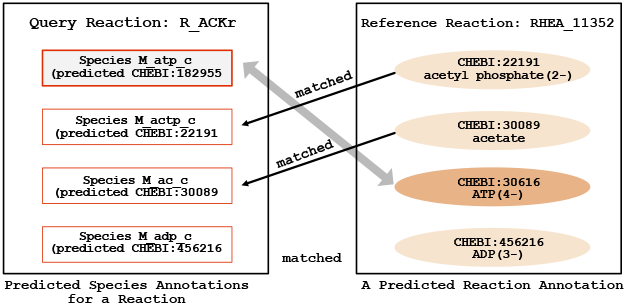
Example of iterative optimization of predicted annotations. The left hand side depicts a reaction for an E. Coli model in BiGG. The right hand side depicts RHEA:11352, the reference reaction with the largest match score for the query reaction. Participants of the query reaction are paired with a participant of the reference reaction. A pairing is “matched” if the annotation of a species in the query reaction is the same as the annotation of its paired species. Predictions can often be improved if the predicted annotation for an unmatched reference species is changed to the annotation of its paired species in the reference reaction.

Actually, we can go further. Once we revise the species annotations based on reaction annotations, we can then revise reaction annotations based on the revisions to species annotations. We provide complete algorithmic details for these revisions in supplemental material S6.

## 3. Results

This section evaluates the effectiveness and efficiency of the AMAS approach to predicting annotations, as well as some details on how one can use AMAS to annotate SBML models.

### 3.1. Evaluations

We evaluate AMAS annotation predictions in terms of effectiveness and computational efficiency. Throughout, our analysis uses MSSC Top with a match score cutoff of 0.0. Our approach compares AMAS annotation predictions with existing annotations in two repositories of biological models, BioModels and BiGG. That is, we assume that the existing annotations are correct (although this may not be the case).

We note that our notions of prediction quality is similar to quality concerns in the field of information retrieval (IR) (e.g., Manning, Raghavan, and Schütze 2008). In IR, there is a set of desired documents (*D*_*D*_) that should be retrieved and there is a set of retrieved documents (*D*_*R*_) that are retrieved. Two key quality measures are recall, 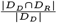, and precision, 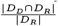, where |*D*| is the size of the set *D*. Unfortunately, these measures have two shortcomings for quantifying AMAS prediction quality. First, recall and precision are correlated because they have the same numerator; this complicates the interpretation of quality measures. Second, AMAS produces an empty prediction set if there is insufficient information to make a prediction. In these cases, precision is undefined since its denominator is 0.

That AMAS intentionally produces empty prediction sets motivates a new metric, *nonempty*. It also is the reason why *accuracy* and *exactness* are only defined if the prediction set is not empty.

To formalize nonempty, accuracy, exactness, we denote the prediction set by *P* and the expected (correct) set of annotations by *E*. First, for a given prediction, *nonempty* is either true or false (1 or 0). Next, if *nonempty* is false, accuracy and exactness are undefined. Otherwise (*nonempty* = 1), we can compute accuracy and exactness:

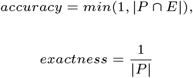

For a recommender system, our feeling is that a list of 3 to 5 choices is acceptable. This means that exactness should be at least 0.2 or 0.33. Note that all three metrics have values in the interval [0, 1], where higher numbers are better.

Tab. 1 summarizes the models we study for predicting species and reaction annotations. Of the 1,000 models in BioModels, only 306 have at least one ChEBI annotated species, and 131 have at least one KEGG or EC-number annotated reaction that can be mapped to Rhea. In BiGG, the fraction of ChEBI and RHEA annotated elements is much higher.

**Table 1.**
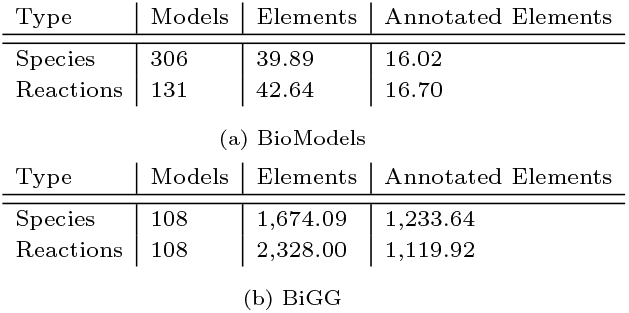
Summary of curated models that contain existing ChEBI and Rhea annotations from 1,000 models in BioModels and 108 models in BiGG. Models are considered only if they have at least one annotation, either species (ChEBI) or reaction (KEGG/EC-number). For these models, the column “Elements” is the average number of species (reactions) in the model, and “Annotated Elements” is the average number of species (reactions) that are annotated.

Tab. 2 reports statistics on AMAS computational efficiency for predicting annotations. We see that the processing time per species for predicting annotations is modest. For both BioModels and BiGG, predicting reaction annotations takes about ten times longer than predicting species annotations. The longer processing times are in large part due to the fact that, for timing purposes, we predict annotations for each reaction separately and so do not take advantage of species predictions that have been done previously. The longer processing times for BiGG reactions is due to having reactions with more participants; two-participant reactions (1 reactant and 1 product) are common in BioModels, and much less so in BiGG.

**Table 2.**
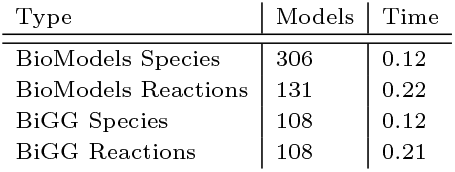
Average processing times (in seconds) for predicting annotations per each species and each reaction in BioModels and BiGG. Reaction times are larger because we include the time to predict annotations for the participant species in the reaction for which prediction is done. BiGG reaction predictions take longer because on the average they have more participants than BioModels reactions.

| Type | Models | Time |
| --- | --- | --- |
| BioModels Species | 306 | 0.12 |
| BioModels Reactions | 131 | 0.22 |
| BiGG Species | 108 | 0.12 |
| BiGG Reactions | 108 | 0.21 |

Fig. 3 plots the accuracy of AMAS predictions for species and reactions in BiGG and BioModels. Accuracy is averaged across all elements of the same type (species or reactions). The figure is organized as two rows of plots. The top row is BiGG and the bottom row is BioModels; the columns are species and reactions. The plots have the same structure. The horizontal axis is the match score cutoff used for MSSC Cutoff; the vertical axis is accuracy. There are results for five Element Filters based on the length of species names. We see that the accuracy of species predictions increases as we increase the minimum length of species names. This make sense since longer names are generally more meaningful.

**Fig. 3.**
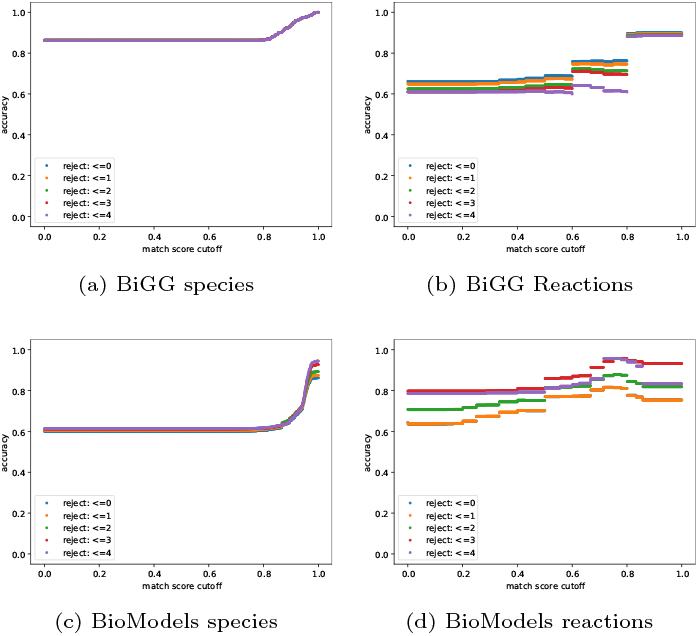
Accuracy of predictions for species (column 1) and reactions (column 2) in BiGG (row 1) and BioModels (row 2). The plots display the relationship between prediction accuracy and the match score obtained by MSSC Top with a cutoff of 0.0. Plots have five lines that reflect element filtering by length of species names (for species) and the number of species in reactions (for reactions).

We also note that accuracy increases with match score cutoff. At first glance, this may seem counter-intuitive since increasing the match score cutoff decreases the size of the prediction set and this in turn can only *decrease* accuracy for a given query element. However, what is happening is that as the match score cutoff increases, *nonempty* decreases. Thus, the query elements for which there are nonempty prediction sets at higher match scores are query elements for which accuracy is larger.

Fig. 4 analyzes AMAS predictions by showing all three quality metrics in combination–*nonempty, accuracy*, and *exactness*. As before, the horizontal axis is the match score cutoff; the particular metrics are indicated by different bar colors. As we saw in Fig. 3, *accuracy* increases with match score cutoff, and this is because *nonempty* is decreasing with match score cutoff. *exactness* also increases with match score cutoff. The quality metrics tend to be better with BiGG than BioModels, which is likely the result of BiGG having longer names for species. We see that *exactness* is always in excess of 0.3, our requirement for a recommender. The accuracy of species predictions tend to be larger than those for predicting reaction annotations. This is expected since reaction predictions depends on the correct prediction of species annotations.

**Fig. 4.**
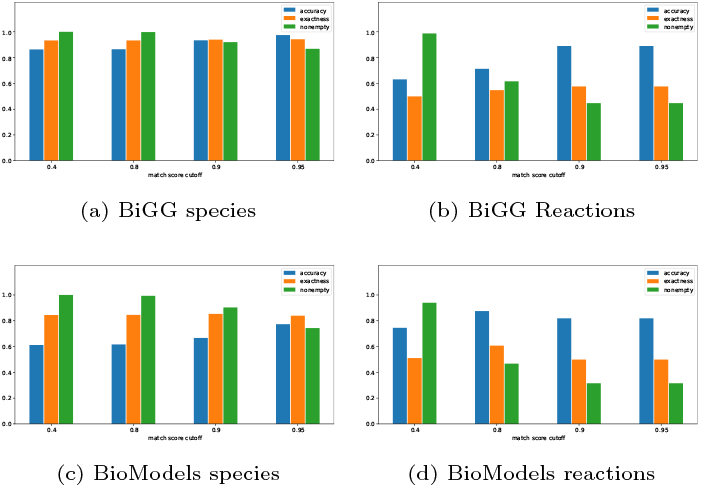
Quality metrics at different match score cutoffs. Element filtering eliminates species names that have less than 3 characters.

### 3.2. Using AMAS

AMAS is an open source, pip-installable Python package. The source code is available at https://github.com/sys-bio/AMAS. The current version was developed and tested on Python version 3.11.2.

AMAS can be used on the command line to recommend and apply annotations using a specified match score cutoff. Fig. 5 shows an example of getting recommendations for all species and reactions in a model, with a match score cutoff of 0.9. Fig. 5 also shows AMAS saving those recommendations as annotations in a new SBML model file. AMAS has multiple scripts that can be run at the command line. Full details can be found in readthedocs at https://amas.readthedocs.io/en/latest/index.html.

**Fig. 5.**
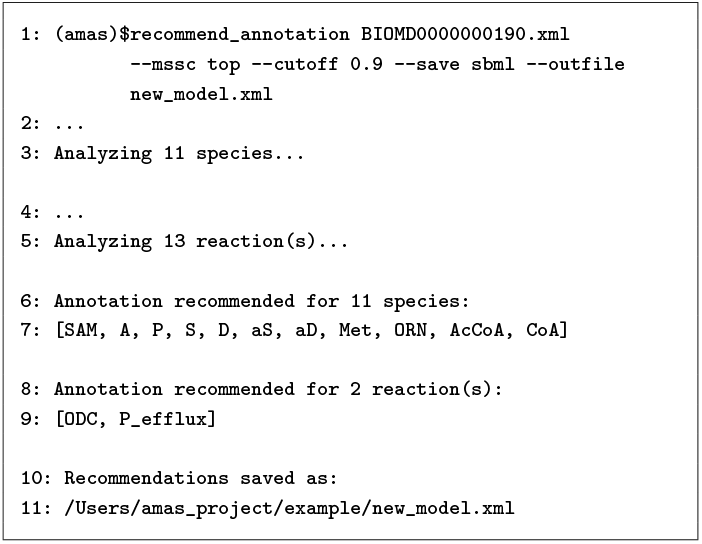
Example of running AMAS on the command line. Line numbers are included to facilitate references to the figure. In the invocation of recommend_annotation (line 1), the only required argument is the path of the SBML model file. Recommendations are saved for model elements with match scores at or above the cutoff (defaults to 0). There are 13 reactions in the model (line 5), but recommendations are mode for only two reactions (lines 8, 9). Recommendations are made for all 11 species (lines 6, 7). The number of recommended reactions will increase if the user applies a smaller cutoff. Finally, annotations in the model were updated using recommended annotations and saved to a new file (lines 10, 11).

Fig. 4 provides guidance as to how to specify the match score cutoff in the command line invocation of AMAS. The user should start by specifying a larger match score cutoff, say 0.9 so that accuracy is larger. If *nonempty* is too small (as indicated by the number of annotations recommended by the command line tool), the user should reduce the match score cutoff. Since accuracy decreases with the match score cutoff, the user should review carefully annotations that are recommended at a lower match score cutoff.

## 4. Discussion and Future Work

High quality annotations provide researchers with a critical understanding of the meaning and behavior of biomedical models. These insights increase the model’s utility and increase the likelihood of others extending or otherwise reusing the model. However, providing high-quality annotations is time consuming and knowledge intensive. As a result, it is common that models have missing or incorrect annotations.

The contributions of this paper are:

1. *A general framework for predicting annotations of model elements*. The framework, shown in Fig. 1, makes use of an existing database of annotated model elements to predict a set of annotations. We specify three metrics for evaluating the quality of prediction: *nonempty, accuracy*, and *exactness*. The framework is instantiated by specifying a match score function for the type of element whose annotations are being predicted.
2. *Match score functions for predicting annotations for species and reactions*. These match score functions are simple to implement and computationally efficient.
3. *An evaluation of prediction efficiency and quality*. We evaluated our approach for predicted annotations for species and reactions in BiGG and BioModels.
4. *AMAS, an open-source package for recommending SBML annotations for species and reactions*. The package is pip-installable.

Our evaluations show that, for SBML models in BiGG and BioModels, AMAS can efficiently and effectively be used to recommend annotations. With these models, AMAS consistently has an *exactness* in excess of 0.3, which means that the prediction set has about 3 elements. Managing the accuracy of AMS prediction requires some consideration of the match score cutoff, the minimum match score at which annotations are predicted. We see that prediction accuracy exceeds 90% in BiGG if the match score cutoff is at least 0.9; similar prediction accuracies are achieved for BioModels reactions. Accuracy is a bit lower in BioModels species, about 0.8 with a 0.95 match score cutoff. This is likely because species names in BioModels are much shorter than in BiGG (averaging 7.2 characters in BioModels versus 19.3 characters in Bigg). A larger match score cutoff reduces *nonempty*, which means that fewer predictions are made. Users should first set a high cutoff to obtain high accuracy predictions. Then, the match score cutoff can be reduced to increase *nonempty*. Greater scrutiny should be applied to predictions made with a lower match score cutoff.

Our near-term efforts will extend the annotation capabilities of AMAS from small molecules in metabolic pathways to larger molecules (e.g., proteins, mRNA, DNA) that are frequently part of signaling pathways. Another near-term direction is to annotate SBML elements beyond species and reactions, such as SBML elements for *model* and *kinetic law*. Reactome and SABIO-RK (Gillespie et al. 2022; Wittig et al. 2018) are possible reference databases to predict annotations for these elements.

Although AMAS can be used as a command-line tool today, we plan to create an API that supports the full AMAS functionality. Such an API would allow for embedding annotation prediction within other software, such as GUI-based or web-based applications. Since our overall goal is to reduce the effort needed to create annotations, any annotation prediction capability should be embedded in an easy-to-use application that model developers might already be using. For example, if a modeler is using software to design or build the model, then if that software calls AMAS to make predictions, these can be reviewed immediately by the modeler, and they can select the most appropriate annotation. Such an interface would also allow AMAS to learn user preferences about annotations (such as via user profiles, as is done in product recommender systems).

In addition, we also plan to explore the ability of AMAS to act as a *critiquing* system. In our review of published models (e.g., those in the BioModels repository), we have numerous examples where the annotation or the name of the element (reactions or species) are incorrect, misleading, or ambiguous. With an appropriate user interface, AMAS could compare the match score of the existing annotation versus a *de novo* prediction for that element. This capability would effectively allow for “debugging” of existing annotations, and would help provide a more reusable library of annotated models.

In sum, we have proposed a general framework for predicting annotations of model elements, and a tool, AMAS, that implements these capabilities. As our evaluation of AMAS with SBML models from BiGG and BioModels demonstrates, our system can efficiently produce accurate annotations. We plan to apply these AMAS capabilities both prospectively, by embedding AMAS within model-development software, and retrospectively, by using AMAS to debug existing models. In both situations, the goal is to make models more understandable and reusable by providing quality annotations about model elements.

## 6. Author contributions statement

W.S. developed the software and conducted data analyses, and wrote the initial draft of the manuscript. J.H. designed the analyses. Together, J.H. and J.G. restructured and revised the initial draft. H.S. conceived the project, provided feedback on the software and reviewed the manuscript. All authors reviewed, revised, and approved the final version of the manuscript.

## 7. Acknowledgments

We acknowledge useful discussions with Lucian Smith and Maxwell Neal.

## References

Alcántara, Rafael et al. (2012) “Rhea—a manually curated resource of biochemical reactions”. In: Nucleic Acids Research 40.D1.

Ashburner, Michael et al. (2000) “Gene Ontology: tool for the unification of biology”. In: Nature Genetics 25.1.

Aziz, Ramy K et al. (2008) “The RAST Server: Rapid Annotations using Subsystems Technology”. In: BMC Genomics 9.1.

Consortium, The UniProt (2014) “UniProt: a hub for protein information”. In: Nucleic Acids Research 43.D1.

Courtot, Mélanie et al. (2011) “Controlled vocabularies and semantics in systems biology”. In: Molecular Systems Biology 7.543.

Cowan, Ann E et al. (2019) “ModelBricks—modules for reproducible modeling improving model annotation and provenance”. In: npj Systems Biology and Applications 5.1.

Degtyarenko, Kirill et al. (2008) “ChEBI: a database and ontology for chemical entities of biological interest”. In: Nucleic Acids Research 36.suppl 1.

Dias, Oscar et al. (2015) “Reconstructing genome-scale metabolic models with merlin”. In: Nucleic Acids Research 43.8.

Dräger, Andreas et al. (2015) “SBMLsqueezer 2: context-sensitive creation of kinetic equations in biochemical networks”. In: BMC Systems Biology 9.1.

Finn, Robert D et al. (2011) “HMMER web server: interactive sequence similarity searching”. In: Nucleic Acids Research 39.suppl 2.

Gillespie, Marc et al. (2022) “The reactome pathway knowledgebase 2022”. In: Nucleic Acids Research 50.D1.

Han, Jiawei et al. (2012) “Getting to know your data”. In: Data mining. Vol. 2. Morgan Kaufmann Boston, MA, USA

Henry, Christopher S et al. (2010) “High-throughput generation, optimization and analysis of genome-scale metabolic models”. In: Nature Biotechnology 28.9.

Krause, Falko et al. (2010) “Annotation and merging of SBML models with semanticSBML” in: Bioinformatics 26.3.

Leonidou, Nantia et al. (2023) “SBOannotator: A Python Tool for the Automated Assignment of Systems Biology Ontology Terms”. In: preprint.org.

Leray, Matthieu et al. (2019) “GenBank is a reliable resource for 21st century biodiversity research”. In: Proceedings of the National Academy of Sciences 116.45.

Manning, Christopher D. et al. (2008) Introduction to Information Retrieval. Cambridge University Press.

McGinnis, Scott and Thomas L Madden (2004) “BLAST: at the core of a powerful and diverse set of sequence analysis tools”. In: Nucleic Acids Research 32.suppl 2.

Mendoza, Sebastián N et al. (2019) “A systematic assessment of current genome-scale metabolic reconstruction tools”. In: Genome Biology 20.1.

Mistry, Jaina et al. (2021) “Pfam: The protein families database in 2021”. In: Nucleic Acids Research 49.D1.

Neal, Maxwell Lewis et al. (2019) “Harmonizing semantic annotations for computational models in biology”. eng. In: Briefings in bioinformatics 20.2.

Needleman, S. B. and C. D. Wunsch (1970) “A general method applicable to the search for similarities in the amino acid sequence of two proteins”. In: J. Mol. Biol. 48.

Nursimulu, Nirvana et al. (2022) “Architect: A tool for aiding the reconstruction of high-quality metabolic models through improved enzyme annotation”. In: PLOS Computational Biology 18.9.

Pruitt, Kim D et al. (2007) “NCBI reference sequences (RefSeq): a curated non-redundant sequence database of genomes, transcripts and proteins”. In: Nucleic Acids Research 35.suppl 1.

Sarwar, Dewan M. et al. (2019) “Model annotation and discovery with the Physiome Model Repository”. In: BMC Bioinformatics 20.1. (Visited on 04/22/2023)

Schulz, Marvin et al. (2011) “Retrieval, alignment, and clustering of computational models based on semantic annotations”. eng. In: Molecular systems biology 7.

Shin, Woosub et al. (2021) “SBMate: A Framework for Evaluating Quality of Annotations in Systems Biology Models”. In: bioRxiv.

Snoep, Jacky L et al. (2006) “Towards building the silicon cell: A modular approach”. In: Biosystems 83.2.

Wittig, Ulrike et al. (2018) “SABIO-RK: an updated resource for manually curated biochemical reaction kinetics”. In: Nucleic Acids Research 46.D1.

