## Supplementary information for "An Automated Model Annotation System (AMAS) for SBML Models"

#### S1 Expressing ChEBI Identifiers as Chemical Formulas

In the ChEBI database, a single chemical species may have one *primary* identifier and several *secondary* identifiers. For example, as shown in Fig. S1, *water* can be described by CHEBI:15377, its primary identifier, but also by secondary identifiers such as CHEBI:13352 and CHEBI:5585. To solve the possible issue of identifiability, `AMAS` recommends species annotations only in primary identifiers.

We briefly describe considerations that `AMAS` makes for species with multiple annotations and/or multiple chemical formulas. Fig. S1 describes the existing ChEBI annotations of *M_h2o_e* in the `e_coli_core.xml` model. This species was annotated with 20 ChEBI identifiers including 6 primary identifiers, indicating protonation and deprotonation of the molecules. As a result, `AMAS` shortens existing or predicted ChEBI annotations into chemical formulas without hydrogen atoms when evaluating annotations or predicting reaction annotations using them. This would also reduce the computational cost of `AMAS`, as fewer unique molecules would be considered when making predictions. In our study, a total of 148,331 unique ChEBI primary identifiers could be mapped to 49,260 unique chemical formulas; the number was further reduced to 22,600 when the formulas were shortened.

#### S2 Running `AMAS` as a Command-line Tool

`AMAS` has multiple scripts that can be run at the command line. Full details can be found in `readthedocs` at https://amas.readthedocs.io/en/latest/index.html.

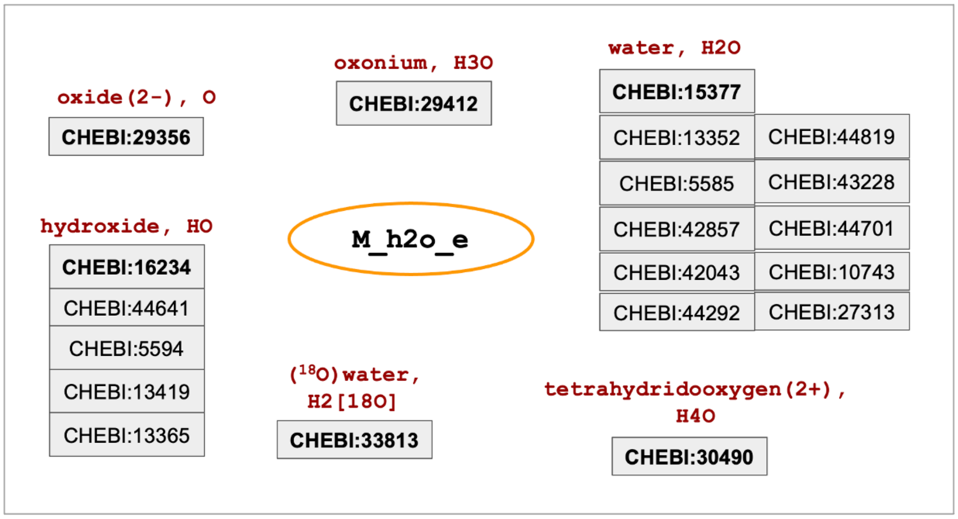


Fig. S1: Existing ChEBI annotations of *M_h2o_e*, a chemical species in the `e_coli_core.xml` model from the BiGG repository. A total of 20 unique identifiers were provided; they could be categorized into 6 unique groups, each with a primary identifier (bold) with a unique chemical formula. Even though the ID indicates *water (H2O)*, additional identifiers indicating protonation and deprotonation were also included in the existing annotations of the model.

### S3 Statistics Related to `AMAS` Predictions

This section provides more detailed statistics about `AMAS` predictions.

From S2a, we see the effect of filtering on the length of species names. The filters we consider result in an empty prediction set for query names that are shorter than 4 characters. But from the histograms, we see that over 98% of the species names in BiGG and over 95% of the species in BioModels have a length of at least 4 characters. This suggests that there is only a modest decrease in *nonempty* if we filter query strings shorter than 4 characters.

S3 analyzes the match score cutoff in BiGG and BioModels for species and reactions. Recall that the match score cutoff is used by `AMAS` to filter the prediction set; an annotation is in the prediction set only if the match score for that annotation is at least as large as the match score cutoff. In BiGG, a match score cutoff of 0.6 results in *nonempty* being over 90% for both species and reactions. In BioModels with a match score cutoff of 0.6, *nonempty* is very large for species, almost 100%, and for reactions *nonempty* is over 70%. Of course, we know from the main text that prediction accuracy is larger with a larger match score cutoff.

These statistics motivate the iterative approach to annotations that we advocate

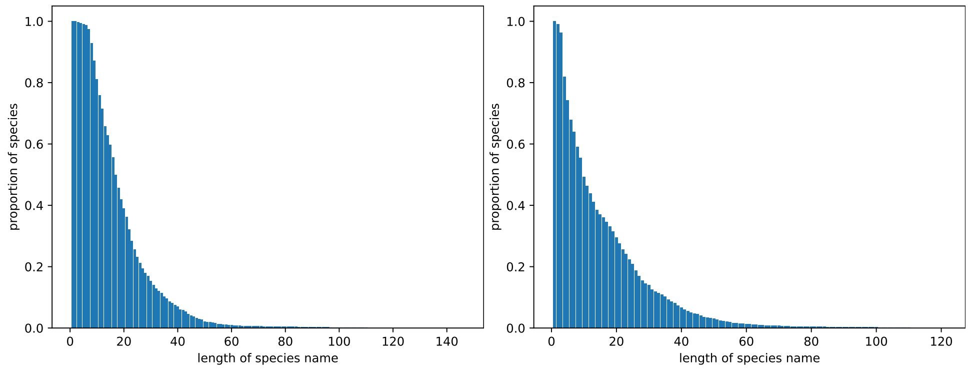


(a) BiGG (b) BioModels

Fig. S2: Bar plots of name lengths.

in the main text. That is, the user should start with a large match score cutoff, say 0.9. If *nonempty* is too low, then decrease the match score, but scrutinize the additional annotations that are predicted.

Fig. S4 displays details of *exactness* for different name length filters for species and reactions in BiGG and BioModels. As expected, *exactness* increases with the minimum match score (match score cutoff). Also note, that even for very small match scores with an element filter of 0, on the average *exactness* is 0.4, which means that the size of the prediction set averages a bit over two annotations.

### S4 Levenshtein Distance

This section provides more details about the use of Levenshtein distance in `AMAS`.

At first glance, the string matching for `AMAS` match scores for species names sounds like polymer matching as is DNA, RNA, and proteins. However, the Needleman–Wunsch algorithm and related approaches assume much longer strings, and work best with a small alphabet and non-binary scoring for character mismatches. This led us to consider the Levenshtein distance, a calculation of the minimal number of insertions, deletions and replacements to make the two strings equal. We define the match score between the query species $Q$ with string $S$ and the reference species $T$ with synonyms $S_1, \cdots, S_n$ as:

$$eScore(Q,T) = 1.0 - \min_{1 \leq j \leq n} (1.0, \frac{d(S, S_j)}{\max(|S|, |S_j|)}),$$

where $d(S, S_j)$ is the Levenshtein distance between the strings $S$ and $S_j$. $|S|$ and $|S_j|$ represent the lengths of the strings.

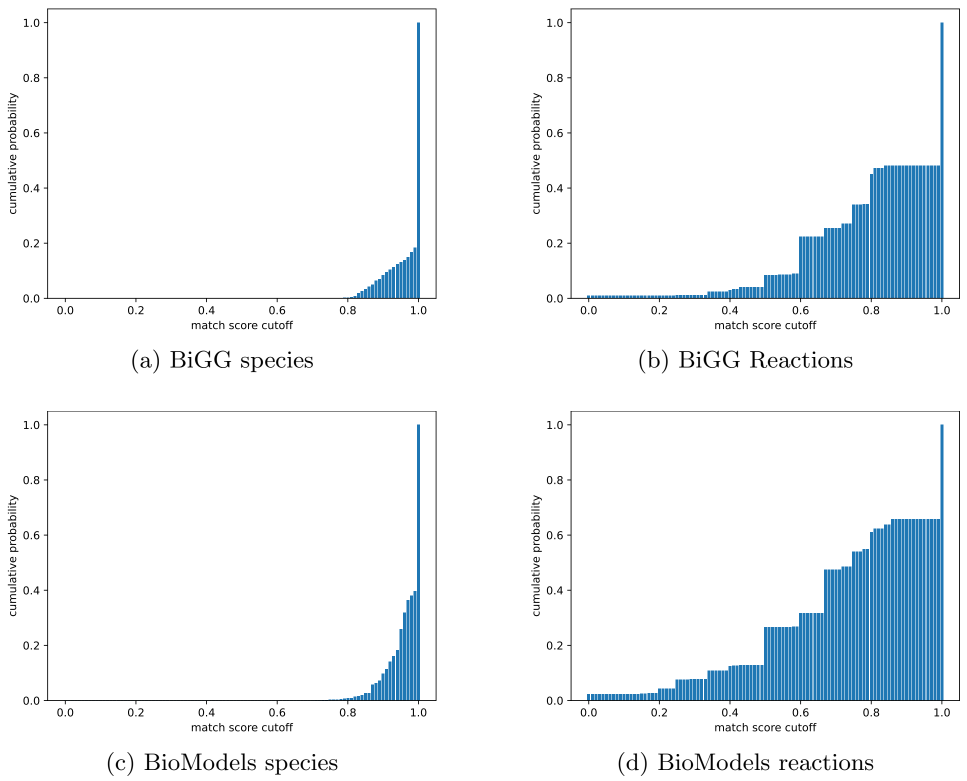


(a) BiGG species

(b) BiGG Reactions

(c) BioModels species

(d) BioModels reactions

Fig. S3: Match score CDF.

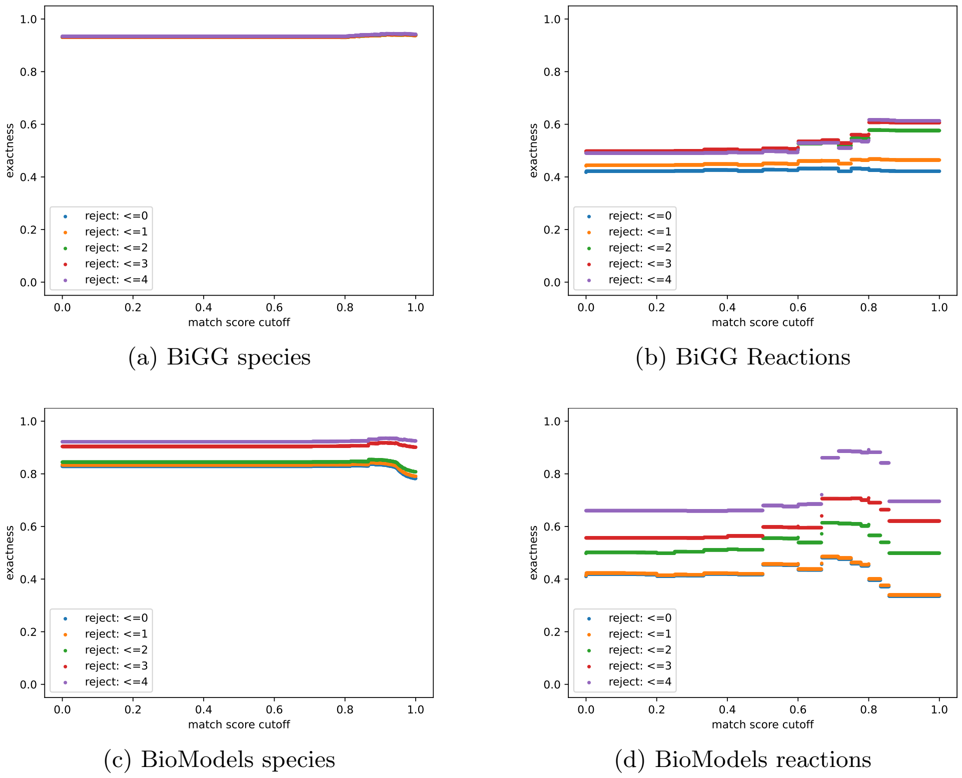


(a) BiGG species

(b) BiGG Reactions

(c) BioModels species

(d) BioModels reactions

Fig. S4: Exactness.

We illustrate the formula with a fsimple example. Suppose that the query species has $S =$ "d_glucose" and that $T =$ CHEBI:17634. This ChEBI term has a single synonym "d-glucose", and so $S_1 =$"d-glucose". Note that $d(S, S_1) = 1$, since $S$ and $S_1$ differ by a single character, and $\max(|S|, |S_1|) = 9$ as both have 9 characters. So, $eScore(Q, T) = 1.0 - min(1.0, \frac{1}{9}) \approx 0.89$.

*eScore* provides good accuracy for predicting annotations, but it is computationally intensive because of calculating Levenshtein distances. This is a serious problem since the computational complexity of predicting annotations is the computational complexity of calcuating the string match score multiplied by the number of reference species (over 140,000) multiplied by the number of synonyms for reference species (up to 74 for each species).

### S5: Calculating rScores

In our analysis of reaction match scores, we found that the number of shared species between the query reaction and the reference reaction was the most important factor to make reasonable predictions. In the current rScore formula, the numerator $(R^q \cdot R^r)$ represents this: the number of matches between the query and the reference elements. We tried a few other options for normalizing the numerator, such as the number species in reference reactions or that in the query reaction, but we found that overall accuracy, exactness, and nonempty to be the highest with the current method.

Here, we provide a detailed example of calculating the rScore. Suppose we have the following reactions:

$$R^q : X \rightarrow Y$$

$$R^{r1} : X + Y \rightarrow Z$$

$$R^{r2} : X + Y \rightarrow W + U$$

$$R^{r3} : Y \rightarrow V$$

where $X, Y, Z, W, U, V$ represent different species. According to the algorithm, (1) the first step is to find the maximum number of matches ($M^q$) between the query reaction $R^q$ and the reference reactions $R^{r1}$, $R^{r2}$, and $R^{r3}$. Since $R^{r1}$ and $R^{r2}$ contain both $X$ and $Y$, $M^q = 2$. (2) Next, the subset of reference reactions, $\mathcal{L}^q$, that have $M^q$ (i.e., 2) matches with the query reaction is $\{R^{r1}, R^{r2}\}$. (3) Now, $D$ is the smallest number in $\{R^{r_j} \cdot R^{r_j} | R^{r_j} \in \mathcal{L}^q\}$, so $D = R^{r1} \cdot R^{r1} = 3$. Then, the rScores are computed as below (rounded to two decimal places):

$$rScore_{R^{r1}} = \frac{2}{3} = 0.67$$

$$rScore_{R^{r2}} = \frac{2}{3} = 0.67$$

$$rScore_{R^{r3}} = \frac{1}{3} = 0.33$$

One alternative method we have considered is Jaccard Index, which is to divide the cardinality of intersection of the query element and the reference element by that of their union. The rScore can be computed using the formula:

$$rScore(R^q, R^r) = \frac{|R^q \cap R^r|}{|R^q \cup R^r|}$$

Therefore, the rScores for the above reference reactions are calculated as below:

$$rScore_{R^{r1}} = \frac{2}{3} = 0.67$$

$$rScore_{R^{r2}} = \frac{2}{4} = 0.50$$

$$rScore_{R^{r3}} = \frac{1}{3} = 0.33$$

Tab. S1 compares the results obtained by the current method and that by Jaccard Index. We found that Jaccard Index-based rScores produced overall higher *accuracy* and *exactness* than the current method but also yielded significantly smaller value of *nonempty*.

| Method | Accuracy | Exactness | Nonempty |
|---|---|---|---|
| Current | 0.77 | 0.46 | 0.73 |
| Jaccard | 0.78 | 0.69 | 0.31 |

(a) BioModels

| Method | Accuracy | Exactness | Nonempty |
|---|---|---|---|
| Current | 0.69 | 0.43 | 0.90 |
| Jaccard | 0.89 | 0.69 | 0.36 |

(b) BiGG

Table S1: Mean accuracy, exactness, and nonempty obtained by the two different rScore methods. We applied MSSC Top with match score cutoff of 0.6.

### S6: Optimizing Annotations

Predicted annotations can be further improved by exploiting the relationships between reactions and species, as depicted in Fig. 2.

Here, we provide more details on how `AMAS` optimizes predicted annotations. This is done in an iterative manner with `AMAS` updating species annotations based on reaction annotations. And then, `AMAS` updates reaction annotations based on the updated species annotations. The iteration continues until there is no improvement in the quality of the prediction. Prediction quality is qualified by the match score averaged over $rScore$ using updated species annotations. The details are:

1. For each query reaction in the model
   (a) Predict annotations for the query reaction using MSSC Top, and select an annotation.
   (b) Find the reference reaction for the predicted annotation.
   (c) Match the participants of the reference reaction with the participants of the query reaction.
   (d) Use the chemical formulas of the unmatched participants to find likely matches. Construct a list of pairings of a participant in the reference reaction with a participant in the query reaction.
   (e) For each pair, assign the annotation of the participant of the reference reaction to the query participant.
2. Reaction annotations are predicted again. Updated annotations for species are accepted if the resulting rScores increase.
3. If the updates are rejected or the match score does not increase or the iteration count is too large, stop. Otherwise, go to (1).

### S7: Details of Evaluation of `AMAS`

There are some details as to how we extract annotations, especially in BioModels. For species, we used the ChEBI annotations under either *bqbiol:is* or *bqbiol:isVersionOf* qualifiers. Reactions are more complicated because BioModels provides few Rhea annotations. Rather, reactions are more frequently annotated with KEGG.Reaction or EC-number. We addressed this by mapping KEGG.Reaction and EC Number into Rhea terms using the cross-referencing information provided in the Rhea database.
